# LIMBARE: an Advanced Linear Mixed-effects Breakpoint Analysis with Robust Estimation Method with Applications to Longitudinal Ophthalmic Studies

**DOI:** 10.1101/2023.01.27.525884

**Authors:** TingFang Lee, Joel S Schuman, Maria de los Angeles Ramos Cadena, Yan Zhang, Gadi Wollstein, Jiyuan Hu

## Abstract

**Purpose:** Broken stick analysis is a widely used approach for detecting unknown breakpoints where association between measurements is non-linear. We propose LIMBARE, an advanced **li**near **m**ixed-effects **b**reakpoint **a**nalysis with **r**obust **e**stimation, especially designed for longitudinal ophthalmic studies. LIMBARE accommodates repeated measurements from both eyes and overtime, and effectively address the presence of outliers.

**Methods:** The model setup of LIMBARE and computing algorithm for point and confidence interval estimates of the breakpoint was introduced. The performance of LIMBARE and other competing methods was assessed via comprehensive simulation studies and application to a longitudinal ophthalmic study with 216 eyes (145 subjects) followed for an average of 3.7±1.3 years to examine the longitudinal association between structural and functional measurements.

**Results:** In simulation studies, LIMBARE showed the smallest bias and mean squared error (MSE) for estimating the breakpoint, with empirical coverage probability of corresponding CI estimate closest to the nominal level for scenarios with and without outlier data points. In the application to the longitudinal ophthalmic study, LIMBARE detected two breakpoints between visual field mean deviation (MD) and retinal nerve fiber layer thickness (RNFL) and one breakpoint between MD and cup to disc ratio (CDR), while the cross-sectional analysis approach only detected one and none, respectively.

**Conclusions:** LIMBARE enhances breakpoint estimation accuracy in longitudinal ophthalmic studies, while cross-sectional analysis approach is not recommended for future studies.

**Translational Relevance:** Our proposed method and companion software R package provides a valuable computational tool for advancing longitudinal ophthalmology research and exploring the association relationships between ophthalmic variables.

## 1 Introduction

Broken stick analysis is a powerful analytical tool for investigating piecewise-linear association between continuous variables, particularly in cases where non-linear associations are present. The goal of the analysis is to accurately detect breakpoints at which the effect size of the association changes, allowing for piecewise-linear association inference to be made before and after these breakpoints. In ophthalmology, broken stick analysis has been used to study the association between structural and functional measurements in glaucoma^1-3^, where varying levels of association between structural measurements, such as average RNFL thickness, and functional measurements obtained with visual field tests^4-9^ have been observed at different stages of disease severity. These differences may be due to the “floor effect” of structural measurements beyond which no further structural damage can be measured, leading to changes in structural-function association^10^.

Current statistical approaches for broken stick analysis in ophthalmology mainly employ cross-sectional approaches where observations from one eye per subject from a single visit are included in the analysis^9^. The approach negates the possibility of maximizing available data by using information obtained from both eyes of the same subject and evaluating longitudinal processes such as disease trajectory. Alternatively, aggregate of measurements across all visits from both eyes are used without considering the inter-eye correlation and within-subject correlation between repeated measurements^8^. Without considering the correlations of repeated measurements from two eyes within a subject, the breakpoint estimation may be misleading.

To address these shortfalls, we propose a new and advanced linear mixed-effects breakpoint analysis with robust estimation (LIMBARE) that can provide accurate broken stick analysis for ophthalmic studies that assess association relationship between measurements longitudinally sampled from both eyes. Specifically, LIMBARE is built upon the segmented mixed model, which accounts for repeated measurements overtime and between eyes, and additional confounders commonly confronted in human studies^11, 12^. LIMBARE further incorporates the least trimmed squares (LTS) strategy^13, 14^ to mitigate the impact of outliers and provide a robust and accurate point estimate of breakpoints. We also propose a new standard error estimator for LIMBARE to improve the coverage probability of the confidence interval estimate. With LIMBARE (method workflow illustrated in Figure 1), the point and confidence interval (CI) estimates of breakpoints, together with the association effect sizes quantified as slopes of the piecewise linear mixed effects model are reported.

**Figure 1.**
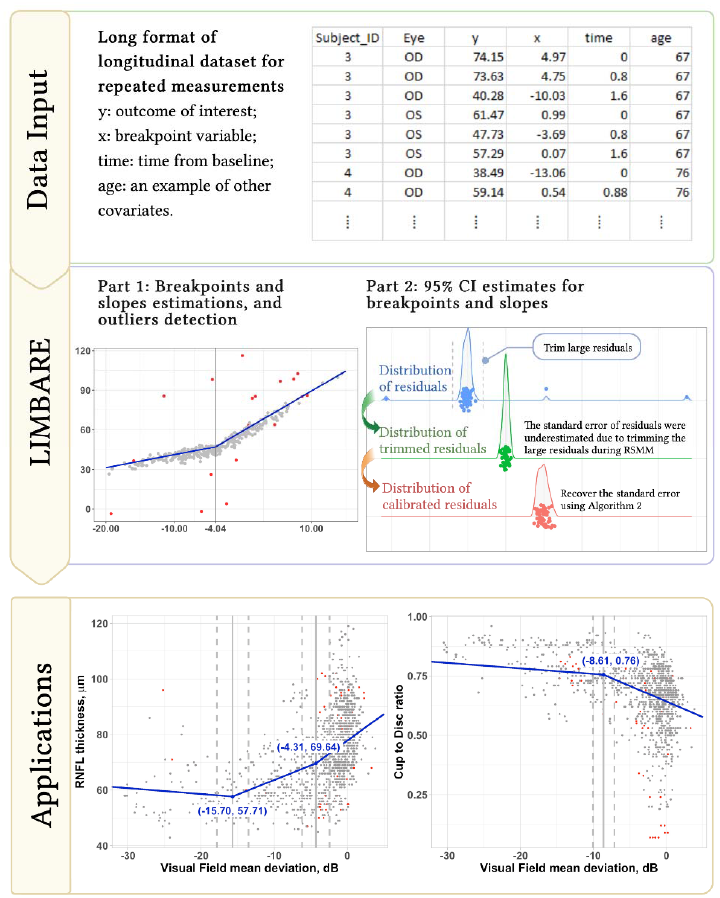
Schematic framework of the linear mixed-effects breakpoint analysis with robust estimation (LIMBARE) method. Data input: the input data is in long format and includes values of the variables of interest (x, y), time (t) and other confounders such as age. LIMBARE: The LIMBARE comprises two parts: Algorithms A1 and A2 are used in each part to detect outliers and obtain robust point (Part 1) and 95% confidence interval (CI) (Part 2) estimates of breakpoints, association slopes, and other unknown parameters. Applications: the output is the piecewise association inference between longitudinal variables, which can be applied to identify changes in association patterns for the study of interested variables.

A simulation study was conducted to evaluate the performance of LIMBARE against competing methods. We also applied these methods to a longitudinal study examining the association between visual field mean deviation (MD) and OCT RNFL thickness, and between MD and cup to disc ratio, to examine the piecewise association relationship. Our results demonstrate that LIMBARE improves the accuracy of breakpoint estimation, and we recommend the use of this method in future ophthalmic studies.

## 2 Methods

### 2.1 Model specification of LIMBARE

We used a segmented mixed-effects model (SMM) to model the longitudinal, non-linear association between two variables of interest. Following the notation of Muggeo (2003)^11^, two variables, denoted by *x*_ijk_ and *y*_ijk_, are observed from subject i (= 1,…, *n*) at visit time *t*_*ijk*_ (*j*= 1, …, *n*_i_) for eye *k* (= “OD”, “OS”). The model also includes time-dependent covariates, denotedby *z*_*ijk*_, such as a constant vector 1, visit time *t*_*ijk*_ and other confounders (See Figure 1 for an illustration of the data input). Our primary goal is to detect unknown yet important breakpoints in the association between *x*_*ijk*_ and *y*_*ijk*_. The SMM regression model is given by Equation (1):

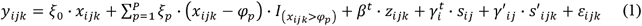

In this equation, Ξ= (ξ 1,-,ξ*p*)^*t*^ is the vector of slope changes between P breakpoints φ_l_,-, φ_P_ (φ_l_ < … < φ_P_), Ψ= (ξ_0_, ξ_0_ + ξ_l_,-, ξ_0_ + ξ_l_ + ξ_2_ +-+ ξ_*P*_)^*t*^ is then the vector of *P*+ 1 slopes (association coefficients) for each of the segments, βis the regression coefficients of the covariates in *z*_*ijk*_ (including intercept), and γ_*i*_ ∼*N*(0, Σ γ) and γ ‘_ij_ ∼N(0, Σ γ ’) are the random effects for covariate *s*_*ij*_ and *s* ‘_*ijk*_ respectively. The residual errors ε_*ijk*_ are assumed to be independent and normally distributed with mean zero and constant variance σ^2^. The model reduces to a standard linear mixed-effects model when no breakpoint is detected (i.e., P= 0).

We estimated the unknown breakpoints Φ= (φ_l_,-, φ_*P*_)^*t*^, and the other unknown parameters 0 = (β^*t*^,ξ0,ξ1, …,ξp, Vech(Σ γ), Vech(Σ γ ’), σ^2^)^*t*^, along with their corresponding 95% confidence intervals (CIs), using the algorithms detailed in **Algorithms 1** and **2** below.

### 2.2 Accurate estimation of breakpoints and other parameters

#### Point estimation

While the specification of the regression model is straightforward, estimating the unknown parameters are challenging due to parameter identifiability problem between Φ and In the case of the standard linear mixed-effects model, maximum likelihood estimation (MLE) approach is commonly used to estimate unknown parameters using the derived log-likelihood function^15^. However, the strategy is not applicable for segmented mixed effects model in Equation (1) because the corresponding log-likelihood function is not differentiable at φ_*p*_ ‘*s*.

Moreover, MLE is sensitive to outliers, which are common in ophthalmic studies due to measurement error, reading error, instrumental error, etc. To overcome these challenges, we propose a robust estimation approach that allows for the detection of multiple breakpoints, accounting for potential outliers.

Given a value 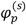, the linear piece (*x*_*ijk*_ - φ_*p*_) · 1(*x*_*ijk*_ > φ_*p*_) can be approximated via the first order Taylor expansion around 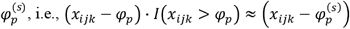, where 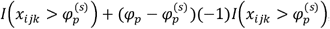 can be considered as the first derivative of (*x*_*ijk*_ - φ_*p*_) · *I*(*x*_*ijk*_ > φ_*p*_). This motivates us to solve the estimation problem by iteratively estimating breakpoints Φ and other parameters Θ. We further introduce the Least Trimmed Squares (LTS) technique to the iteration algorithm to reduce the impact of outliers in the data on parameter estimation^16, 17^.

Specifically, given a predetermined proportion of outlier data points α ∈ (0,1), the iterative procedure for parameter estimation of LIMBARE is given as Algorithm 1 (A1):

##### Algorithm 1

(A1) Point estimate of parameters for LIMBARE framework

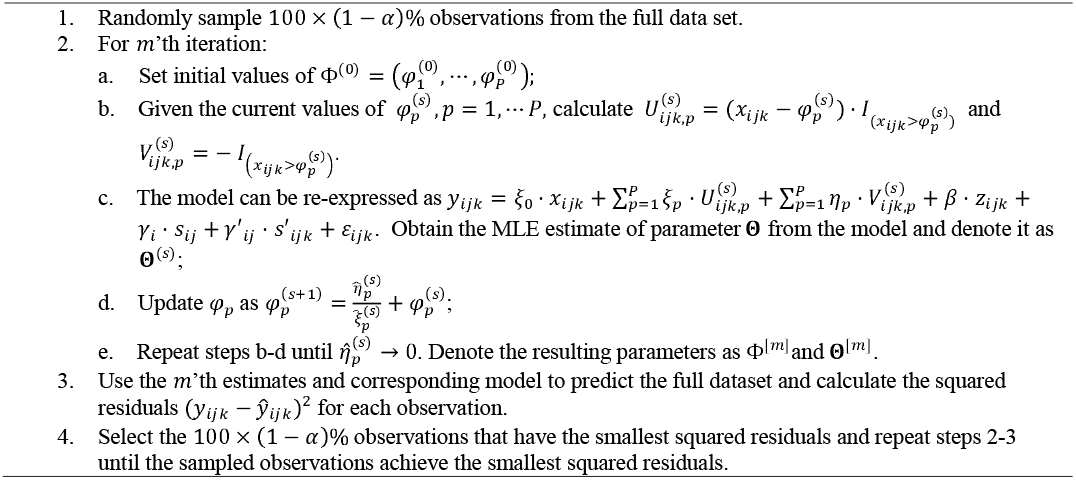

The proposed iterative procedure consists of two layers. The outer layer (Steps 1,3, and 4) employs the LTS technique to account for the potential impact of outliers on the model fitting.

The inter layer (a-e of Step 2) uses the iterative optimization algorithm of the segmented mixed model, which we will discuss later in the competing methods section. To determine the trimming proportion of outliers, we assessed and compared three values of α: 0(no outlier), 0.05, and 0.10 for model fitting and parameter estimation.

#### Estimation of standard error (SE) and confidence interval (CI)

While LTS is effective in reducing or eliminating the influence of outliers on point estimation, it can lead to underestimation of the standard errors of the unknown parameters^18, 19^. To address this issue, we propose Algorithm 2 (A2) to improve the estimates of SEs and CIs of unknown parameters, including breakpoints and their associated slopes.

##### Algorithm 2

(A2) SE and 95% CI estimate of LIMBARE framework

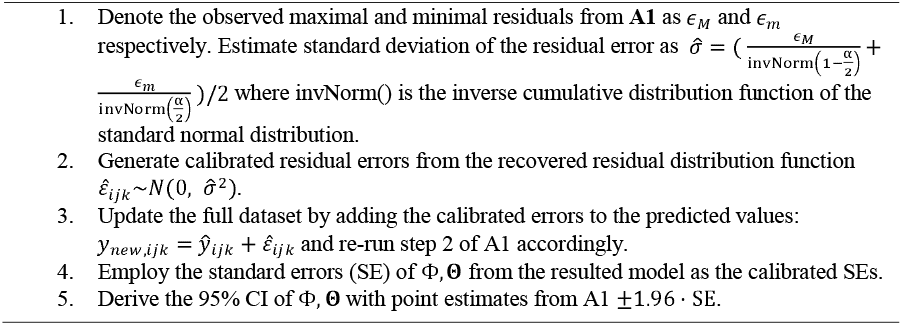

The algorithm approximates the α:/2 × 100% and (1-α/2) x 100% quantiles of the distribution of model residuals with the observed maximal and minimum residuals, since the observed dataset is trimmed with α × 100%. By recovering of distribution of the residuals, each observation ‘s residual is calibrated and used as input for the estimation of SE and CI.

### Hypothesis tests

To test the null hypothesis that the slope of each regression piece is equal to zero, we use Wald test statistics calculated from the point and SE estimates of the slope parameters ‘1= (ξ_0_, ξ_0_ + ξ_l_, …, ξ_0_ + ξ_l_ + ξ_2_ + …+ ξ_*P*_)^*t*^. Specifically, we divide the point estimate of the slope by the corresponding SE to obtain the Wald test statistics. This statistics is then used to examine the null hypothesis for each regression piece.

Figure 1 illustrates the input, algorithm and the output of the proposed method.

### 2.3 Competing methods

To evaluate the performance of our proposed method, we compared it with three alternative methods commonly used for estimating breakpoints in longitudinal studies. The first method is the segmented mixed model (**SMM**) proposed by Muggeo et al^11, 20^. SMM uses an iterative optimization algorithm to estimate the model parameters, but does not account for outlier data points. Therefore, we also evaluated the effect of outliers on the estimates of breakpoints obtained with this method. The second method is the robust mixed-effects segmented model (**RMESR**) proposed by Zhou et. al.(2020)^19^. RMESR combines the estimates from both segmented linear models and linear mixed-effects models, with the breakpoint estimated based on the segmented linear model even though the data comes from a longitudinal design. To accommodate outliers in the dataset, RMESR uses the least trimmed squares technique. The third method is the conventional method, named **CROSS**, in which longitudinal measurements are aggregated cross-sectionally and analyzed with a segmented linear regression model^11^. The detailed description of each method can be found in **Appendix A** of the **Supplementary material**.

### 2.4 Simulation setup

Our simulation was designed to imitate the structural and functional data from real longitudinal ophthalmic studies. For both Scenario 1 (without outliers) and 2 (with outliers), we simulated two breakpoints, φ_l_ and φ_2_, for the variable *x* (referred to as the “breakpoint variable” hereafter). We also included a time variable, t, to record the time from baseline visit in years, and considered random effects between visits of the same subject and between eyes within subjects. To generate the outcome variable y, we used the following model:

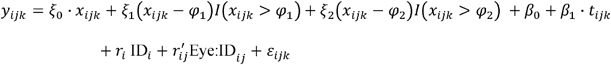

Here, the start slope and slope changes (ξ_0_, ξ_l_, ξ_2_) were set to (1, 2, 3), and the breakpoints (φ_l_, φ_2_) were set to (−10, 2), while the intercept the time slope (p_0_, p_l_) were set to (50, -0.5). We set *x*_*ijk*_ to following a normal distribution *N*(−4, 6^2^), and *t*_*ij*_ to take values (0, 1, 2, 3)^*t*^. The random effects, 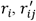, and ε_*ijk*_ were generated from normal distributions with means 0 and variances 1, 4, and 0.25, respectively. We simulated data from 3-4 visits and from two eyes for 60% of the total of 300 subjects. We conducted N = 1000 replications to evaluate the performance of our proposed method, LIMBARE, and compared it with the SMM, RMESR, and CROSS methods.

To simulate outliers under Scenario 2, we randomly selected 5% of observations from the simulated dataset and added (max(*y*) - min (*y)*)/4 to the outcome variable y and (max(*x*) - min (*x*))/4 to the breakpoint variable *x*.

We used the finite sample bias and mean squared error (MSE), and the empirical coverage probability (ECP), as comparison metrics for assessing the point and confidence interval estimates of the model parameters, respectively.

### 2.5 The longitudinal ophthalmic study

This longitudinal clinical study examined 216 eyes (163 open angle glaucoma, 44 glaucoma suspects, and 9 healthy eyes) from a cohort of 145 subjects aiming to investigate changes in ocular structure and function over time. The study was approved by the IRB of the University of Pittsburgh and NYU Langone Health, and all participants provided informed consent^8^. Each subject underwent comprehensive ophthalmic examinations including qualified visual fields, OCT optic nerve head (ONH) and macular scans, in at least 5 visits over an average period of 3.7 years (**Table 1**). We evaluated the piecewise association between visual field mean deviation (MD) and two structural measures: retinal nerve fiber layer (RNFL) thickness and cup to disc ratio (CDR). The proportion of data trimming α was set to 0.05 for the MD-RNFL association, and 0.1 for the MD-CDR association. To control for potential confounding factors, such as time from baseline visit (in year), age at baseline, signal strength of scans, and eye, all break-point analysis models were adjusted accordingly. We used Bayesian information criterion (BIC) to determine the optimal number of breakpoints for each association and each method. The model with the smallest BIC was selected, and corresponding estimated breakpoints, slopes were reported.

**Table 1.** Characteristics of the longitudinal clinical study. This table presents the demographic and clinical characteristics of the study population.

| Parameters | Ns or Mean ± standard deviation (SD) |
| --- | --- |
| Eyes/subjects | 216/145 |
| N visits during f/u | 5.8±0.9 |
| N observations | 1221 |
| Age (years) | 69.7±10.6 |
| follow up (years) | 3.7±1.3 |
| ONH signal strength | 8.3±0.8 |
| baseline RNFL (μm) | 73.8±14.4 |
| baseline MD (dB) | -4.23±6.29 |
| baseline CDR | 0.67±0.15 |

**Table 2:** Simulation results for Scenario 1: synthetic dataset without outliers (breakpoints at -10 and 2). The table shows comparisons of point estimates, standard error estimates, and confidence interval estimates of breakpoints among different statistical methods.

| | $\varphi_1 = -10$ | | | | | $\varphi_2 = 2$ | | | | |
| --- | --- | --- | --- | --- | --- | --- | --- | --- | --- | --- |
|  | Est | Bias | MSE | SE | ECP | Est | Bias | MSE | SE | ECP |
| LIMBARE | -10.001 | -5E-04 | 0.0010 | 0.0291 | 0.923 | 2.0000 | <0.0001 | 0.0005 | 0.0193 | 0.928 |
| SMM | -10.001 | -7E-04 | 0.0008 | 0.0284 | 0.943 | 1.9999 | -0.0001 | 0.0004 | 0.0189 | 0.942 |
| RMESR | -10.038 | -0.038 | 0.0027 | 0.0189 | 0.485 | 2.0340 | 0.0340 | 0.0017 | 0.0126 | 0.349 |
| CROSS | -5.9814 | 4.0186 | 19.7530 | 0.8238 | 0.141 | -1.6171 | -3.6171 | 16.5240 | 0.8236 | 0.141 |

Est, bias and MSE represents the mean value, bias and mean squared error of the point estimate, respectively. SE represents mean value of the standard error estimate of the breakpoint; ECP represents the empirical coverage probability of the 95% confidence interval estimate.

**Table 3:** Simulation results for Scenario 2: synthetic dataset with outliers (breakpoints at -10 and 2). The table shows comparisons of point estimates, standard error estimates, and confidence interval estimates of breakpoints among different statistical methods.

| | $\varphi_1 = -10$ | | | | | $\varphi_2 = 2$ | | | | |
| --- | --- | --- | --- | --- | --- | --- | --- | --- | --- | --- |
|  | Est | Bias | MSE | SE | ECP | Est | Bias | MSE | SE | ECP |
| LIMBARE | -9.9963 | 0.0037 | 0.0817 | 0.0705 | 0.996 | 2.0080 | 0.0080 | 0.1169 | 0.0662 | 0.998 |
| SMM | -10.336 | -0.336 | 7.2465 | 0.8715 | 0.747 | 1.7874 | -0.2126 | 4.8063 | 0.7424 | 0.778 |
| RMESR | -10.073 | -0.073 | 1.2084 | 0.0197 | 0.597 | 1.9793 | -0.0207 | 0.6337 | 0.0139 | 0.567 |
| CROSS | -5.5309 | 4.4691 | 28.4270 | 1.2464 | 0.218 | -1.9612 | -3.9612 | 22.8790 | 1.7822 | 0.231 |

Est, bias and MSE represents the mean value, bias and mean squared error of the point estimate, respectively. SE represents mean value of the standard error estimate of the breakpoint; ECP represents the empirical coverage probability of the 95% confidence interval estimate.

## 3. Results

### 3.1 Simulation results

We present the estimation performance of all assessed methods under two scenarios, including mean value, empirical bias, and MSE for point estimates, mean standard errors (SEs) and empirical coverage probability (ECP) for 95% confident interval estimates of the detected breakpoints in Error! Reference source not found.**2** and Error! Reference source not found.**3**. Additionally, **Table S1** and **Table S2** in **Appendix B** of the **Supplementary material** demonstrate the estimation performance for the estimates of start slope and slope changes for all assessed methods.

Under the first scenario, which involves synthetic datasets without outliers (**Error! Reference source not found.2** and **S1**), the proposed method LIMBARE and SMM are statistically efficient, as indicated by their negligible bias and small MSE for point estimates. In comparison, the point estimate of RMESR and CROSS has a larger bias and MSE than other methods. The CI estimates of LIMBARE and SMM have satisfactory ECPs close to the nominal level of 95%.

However, the CI estimates of RMESR and CROSS tend to have much lower ECPs than nominal. The under-coverage of RMESR is mainly due to the underestimation of SEs, while the poor performance of the CI estimates of CROSS is largely driven by the bias of its point estimate.

Under the second scenario, which involves synthetic datasets with outliers (**Error! Reference source not found.3** and **S2**), the proposed method LIMBARE has the markedly smallest bias and MSE for point estimates, indicating its statistical efficiency and superior robustness to outliers. The point estimates of all competing methods have a large bias and MSE, indicating the failure to accurately detect breakpoints and estimate slopes under this scenario. The ECPs of all competing methods are substantially lower than nominal, while LIMBARE maintains ECP closest to the nominal. However, we detected a slight over-coverage of the CI estimate of LIMBARE.

### 3.2 Broken stick analysis of the longitudinal ophthalmic study

We conducted a piecewise association analysis between visual field mean deviation (MD) and RNFL thickness, as well as MD and CDR, using four different methods: LIMBARE, SMM, RMESR, and CROSS. **Figures 2** and **3**, along with **Tables 4** and **5**, present the results of these analyses.

**Figure 2.**
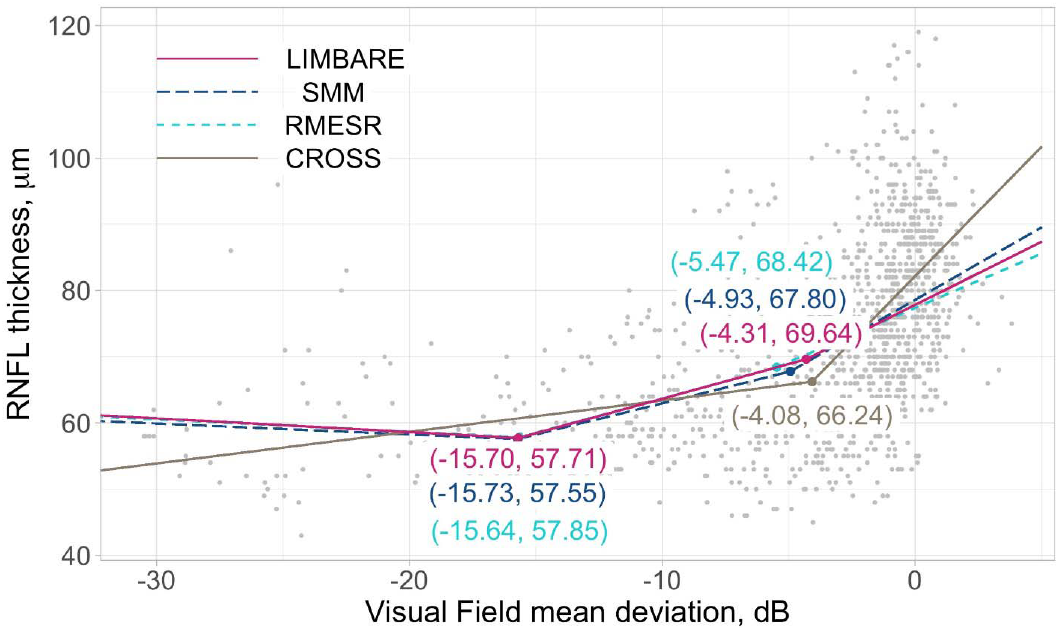
Estimated piecewise associations between visual field mean deviation and RNFL thickness and from all assessed methods.

**Figure 3.**
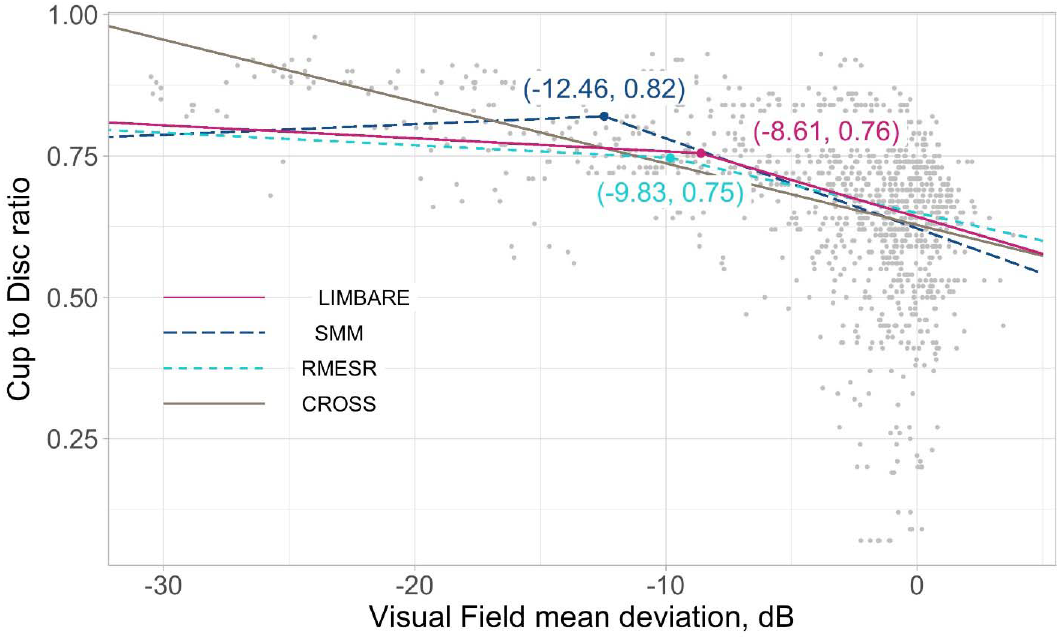
Estimated piecewise associations between visual field mean deviation and cup to disc ratio from all assessed methods.

**Table 4.** Parameter estimates [95% CIs] for the broken stick association analysis between MD and RNFL. Breakpoints are reported as dB and slopes as μm/dB.

|  |  |  |  |  |
| --- | --- | --- | --- | --- |
| LIMBARE | Breakpoints | -15.70<br>[-17.86, -13.53] |  | -4.31<br>[-6.19, -2.42] |
|  | Slopes | -0.21<br>[-0.63, 0.22] | 1.05*<br>[0.81, 1.29] | 1.90*<br>[1.52, 2.29] |
| SMM | Breakpoints | -15.73<br>[-17.87, -13.58] |  | -4.93<br>[-6.25, -3.62] |
|  | Slopes | -0.16<br>[-0.51, 0.18] | 0.95*<br>[0.70, 1.2] | 2.19*<br>[1.90, 2.48] |
| RMESR | Breakpoints | -15.64<br>[-16.95, -14.34] |  | -5.47<br>[-7.16, -3.78] |
|  | Slopes | -0.19<br>[-0.42, 0.05] | 1.04*<br>[0.87, 1.21] | 1.64*<br>[1.40, 1.87] |
| CROSS | Breakpoint | -4.08<br>[-6.26, -1.89] |  |  |
|  | Slopes | 0.48<br>[-0.03, 0.98] |  | 3.91*<br>[1.93, 5.90] |

\* p-value <0.05 for the statistical test against the null hypothesis of slope equal to zero.

**Table 5.** Parameter estimates [95% CIs] for the broken stick association analysis between MD and CDR. Breakpoints are reported as dB and slopes as cup-to-disc-ratio/dB.

|  |  |  |  |
| --- | --- | --- | --- |
| LIMBARE | Breakpoints | -8.61<br>[-10.09, -7.14] |  |
|  | Slopes | -0.0023*<br>[-0.0038, -0.0008] | -0.0131*<br>[-0.0155, -0.0107] |
| SMM | Breakpoint | -12.46<br>[-13.85, -11.07] |  |
|  | Slopes | 0.0019<br>[-0.0011, 0.0048] | -0.0159*<br>[-0.0178, -0.014] |
| RMESR | Breakpoint | -9.83<br>[-10.69, -8.98] |  |
|  | Slopes | -0.0022*<br>[-0.0030, -0.0014] | -0.0099*<br>[-0.0109, -0.0089] |
| CROSS | Slope<br>(breakpoint not detected) | -0.0109*<br>[-0.0141, -0.0077] |  |

\* p-value <0.05 for the statistical test against the null hypothesis of slope equal to zero.

For the MD-RNFL association (**Figure 2** and **Table 4**), LIMBARE, SMM, and RMESR consistently identified two breakpoints, while CROSS detected only one. Specifically, LIMBARE indicated that RNFL thickness decreases significantly and rapidly at a rate of 1.9 [1.52,2.29] μm/dB before reaching the breakpoint of -4.31 dB (from top right to bottom left in Figure 2, which represents the disease spectrum from mild to severe glaucoma). After that, RNFL thickness still significantly decreases but at a slower rate of 1.05 [0.81,1.29] μm/dB until the next breakpoint of -15.70 dB. The association between MD and RNFL approaches a plateau with insignificant rate of change when MD is smaller than -15.70 dB. In contrast, CROSS captured only one breakpoint at MD= -4.08 dB, which is inconsistent with the other three methods. The estimated slope from CROSS indicated that RNFL thickness decreases significantly with MD when MD is larger than -4.08 dB, which is not typically observed clinically. Noticeably, RMESR reported the narrowest confidence interval estimates for both breakpoints and slopes, which suggests the possibility of under-coverage of the ground truth of these parameters (**Table 4**).

For the MD-CRD association (**Figure 3** and **Table 5**), LIMBARE, SMM, and RMESR identified one breakpoint while Cross detected none. LIMBARE and RMESR, which were designed to accommodate outliers, reported breakpoints close to each other at -8.61 dB and -9.83 dB, respectively. The corresponding slope estimates from LIMBARE and RMESER indicated that CDR increases at the rate of 0.0131 [0.0107,0.0155] per dB (from right to left of Figure 3) and 0.0099 [0.0089,0.0109] per dB before MD decreases to the breakpoint. After that, the increasing trend slows down with a rate of 0.0023[0.0008,0.0038] per dB and 0.0022[0.0014,0.0030] per dB. Yet, RMESR reported much narrower confidence interval estimates for both breakpoints and slopes. In comparison, SMM showed that CDR increases at the rate 0.0159 [0.0140,0.0178] per dB when MD decreases before reaching the breakpoint of -12.46 dB. After the breakpoint, CDR decreases at the rate 0.0019 [-0.0011,0.0048] per dB as MD declines. The patterns identified by LIMBARE and RMESR complied with clinical observation, while SMM showed a trend that is less frequently observed. RMESR resulted the narrowest confidence interval estimates compared with LIMBARE.

Overall, the results suggest that LIMBARE is a robust method for identifying breakpoints in the association between MD and RNFL thickness or MD and CDR by providing both point and confidence interval estimates. Conventional methods like CROSS may miss these breakpoints, leading to inaccurate or incomplete understanding of the relationship between these variables. The identified breakpoints and slopes may provide valuable clinical insights into the study of glaucoma.

## 4 Conclusion

In this manuscript, we proposed a novel advanced linear mixed-effects breakpoint analysis with robust estimation (LIMBARE) for longitudinal data analysis. The proposed method provides efficient point estimation of breakpoints and other association parameters, largely mitigating the impact of outliers, and improving the estimation of confidence intervals for both breakpoints and slopes. Simulation studies show that LIMBARE outperforms three competing methods in terms of negligible bias, smallest mean squared error, and empirical coverage probabilities closest to the nominal level of 95% under the scenarios with and without outliers.

In the application to broken stick analysis to a longitudinal clinical study, the piecewise associations between MD and RNFL thickness estimated by LIMBARE, SMM, and RMESR were comparable and consistent with clinical observations. The piecewise associations between MD and CDR estimated by LIMBARE and RMESR also complied with clinical observation, while SMM exhibited a less likely clinical pattern. The results of CROSS, however, were inconsistent with three other methods.

In conclusion, LIMBARE is recommended for future longitudinal ophthalmic studies as it exhibited the most favorable results in both simulation scenarios and illustrated patterns that comply with clinical observation. Furthermore, a user-friendly R package “LIMBARE” is publicly available at https://github.com/JiyuanHu/. The conventional cross-sectional method is not capable of capturing breakpoints in longitudinal studies and its use is not recommended.

