## Supplementary material for "LIMBARE: an Advanced Linear Mixed-effects Breakpoint Analysis with Robust Estimation Method with Applications to Longitudinal Ophthalmic Studies"

### **Appendix A: Optimization algorithms of competing methods**

**1. CROSS**

One practical approach for conducting breakpoint association analysis in glaucoma research is to aggregate repeated measurements within each subject by taking the means and estimate breakpoints cross-sectionally with a segmented linear regression model (Muggeo 2003). This method, referred to as CROSS in this manuscript, involves calculating the mean values of the breakpoint variable, response variable, and covariates averaged over time, denoted by $\bar{x}_{i}$, $\bar{y}_{i}$ and $\bar{z}_{i}$, respectively. The segmented linear model is expressed as

$\bar{y}_{i}=\xi_{0}\cdot\bar{x}_{i}+\sum_{p=1}^{P} \xi_{p}\cdot\left( \bar{x}_{i}-\varphi_{p} \right)\cdot I_{\left( \bar{x}_{i}-\varphi_{p} \right)}+\beta^{t}\cdot\bar{z}_{i}+\varepsilon_{i}$,

where $\varepsilon_{i}\sim N(0, \sigma^{2})$. The parameter notations are the same as those used in LIMBARE without causing confusion. The parameter estimates can be obtained using the iteration algorithm A3 as follows:

**Algorithm 3 (A3) Parameter Estimation for Segmented Linear Regression Model (CROSS)**

1. Set the initial value of $\Phi^{(0)}=\left( \varphi_{1}^{\left( 0 \right)},\cdots, \varphi_{P}^{\left( 0 \right)} \right).$

2. Given the current value of $\varphi_{p}^{\left( s \right)}$, calculate $U_{i, p}^{(s)}=(\bar{x}_{i}-\varphi_{p}^{\left( s \right)})\cdot I(\bar{x}_{i}>\varphi_{p}^{\left( s \right)})$ and $V_{i, p}^{(s)}=- I(\bar{x}_{i}>\varphi_{p}^{\left( s \right)})$.

3. Fit the regression model to the aggregated dataset: $\bar{y}_{i}=\xi_{0}\cdot\bar{x}_{i}+\sum_{p=1}^{P} \xi_{p}\cdot U_{i, p}^{\left( s \right)}+\sum_{p=1}^{P} \eta_{k}\cdot V_{i, p}^{\left( s \right)}+\beta^{t}\cdot z_{i}+\varepsilon_{i}$. Obtain the maximum likelihood estimates and corresponding standard error estimates of parameter $\boldsymbol{\Theta}$, and denote it as $\boldsymbol{\Theta}^{(s)}$ and SE($\boldsymbol{\Theta}^{(s)}$) respectively.

4. Update $\varphi_{p}$ with $\varphi_{p}^{\left( s+1 \right)}=\frac{\hat{\delta}_{p}^{(s)}}{\hat{\xi}_{p}^{(s)}}+\varphi_{p}^{\left( s \right)}$ where $\hat{\delta}_{p}^{(s)}=\hat{\xi}_{p}^{(s)}\cdot(\varphi_{p}^{\left( s \right)}-\varphi_{p}^{\left( 0 \right)})$.

5. Repeat steps 2, 3, and 4 until $\hat{\delta}_{p}^{(s)}\to0$.

6. Derive the 95% confidence interval of $\mathrm{parameters}$ with point estimates from step 5 $\pm1.96\cdot\text{SE}$.

**2. Segmented mixed model (SMM)**

Muggeo (2022) proposed the segmented mixed model (SMM) for breakpoint analysis of longitudinal datasets. The model setup used in SMM is the same as Equation (1) in our proposed method LIMBARE. SMM can be viewed as a simplified version of the LIMBARE method, where the proportion of outliers $\alpha$ is fixed at 0. Unlike LIMBARE, SMM only uses a-e of step 2 from Algorithm A1, and as a result, SMM does not account for the potential impact of outliers on parameter estimation.

**3. Robust mixed-effects segmented model (RMESR)**

Zhou et al. 2020 proposed a robust mixed-effects segmented model (RMESR) that combine the parameters estimates of both segmented linear model and linear mixed-effects model. The employ the least trimmed squares (LTS) technique for breakpoint detection, which was robust against outliers in the dataset. However, one caveat of using LTS is that it may underestimate the standard error of parameters, resulting in confidence interval estimates with lower coverage probability than nominal. Given an outlier rate of $\alpha\in(0, 1)$,

**Algorithm 4 (A4) Parameter Estimation for Robust Mixed-effects Segmented Model (RMESR)**

1. Sample $100\times\left( 1-\alpha\right)\%$ observations from the data.

2. Set initial values of $\Phi^{(0)}=\left( \varphi_{1}^{\left( 0 \right)},\cdots, \varphi_{P}^{\left( 0 \right)} \right);$
   a. Given the current values of $\varphi_{p}^{\left( s \right)},p=1,\cdots P,$ fit the linear mixed effects model to the dataset: $y_{ijk}=\xi_{0}\cdot x_{ijk}+\sum_{p=1}^{P} \xi_{p}\cdot\left( x_{ijk}-\varphi_{p}^{\left( s \right)} \right)\cdot I_{\left( x_{ijk}>\varphi_{p} \right)}+\beta\cdot z_{ijk}+\gamma_{i}\cdot s_{ij}+{\gamma^{'}}_{ij}\cdot{s^{'}}_{ijk}+\varepsilon_{ijk}.$ Obtain the MLE estimate of all parameters except $\varphi_{p}^{\left( s \right)}$;
   b. Calculate the predicted random effects ${\hat{\text{reff}}}_{ijk}^{(s)}$ , and let $\hat{y}_{ijk}^{\left( s \right)}=y_{ijk}-{\hat{\text{reff}}}_{ijk}^{\left( s \right)}$.
   c. Estimate the breakpoints $\varphi_{p}^{\left( s+1 \right)}$ using Algorithm A3, the segmented linear model with response variable $\hat{y}_{ijk}^{\left( s \right)}$ and breakpoint variable $x_{ijk}$;
   d. Repeat a-c until convergence.

3. Use the current estimates to predict the full data and calculate the squared residuals ${(y_{ijk}-\hat{y}_{ijk})}^{2}$.

4. Select the top $100\times\left( 1-\alpha\right)\%$ observations that have smallest squared residuals.

5. Repeat steps 2, 3, and 4, until convergence.

### **Appendix B: Supplemental results**

| **Table S1**: Simulation results for Scenario 1: synthetic dataset without outliers. Shown are comparisons of point and CI estimation of slope $\xi_{0}$and change of slopes $\xi_{1}, \xi_{2}$ among different statistical methods. | | | | | |
| --- | --- | --- | --- | --- | --- |
| $\xi_{0}=1$ | | | | | |
|  | Est | Bias | MSE | SE | ECP |
| LIMBARE | 0.9998 | -0.0002 | 0.0002 | 0.0118 | 0.915 |
| SMM | 0.9999 | -0.0001 | 0.0001 | 0.0115 | 0.943 |
| RMESR | 0.9877 | -0.0123 | 0.0004 | 0.0078 | 0.544 |
| CROSS | 1.8986 | 0.8986 | 42.975 | 2.2315 | 0.283 |
| $\xi_{1}=2$ | | | | | |
|  | Est | Bias | MSE | SE | ECP |
| LIMBARE | 2.0001 | 0.0001 | 0.0002 | 0.0127 | 0.929 |
| SMM | 2.0000 | 0.0000 | 0.0002 | 0.0124 | 0.953 |
| RMESR | 2.0132 | 0.0132 | 0.0004 | 0.0083 | 0.546 |
| CROSS | 1.7670 | -0.2330 | 218.67 | 66.861 | 0.469 |
| $\xi_{2}=3$ | | | | | |
|  | Est | Bias | MSE | SE | ECP |
| LIMBARE | 3.0004 | 0.0004 | 0.0002 | 0.0127 | 0.925 |
| SMM | 3.0003 | 0.0003 | 0.0002 | 0.0124 | 0.948 |
| RMESR | 3.0143 | 0.0143 | 0.0004 | 0.0084 | 0.526 |
| CROSS | 0.7663 | -2.2337 | 205.53 | 65.599 | 0.321 |

Est, bias and MSE represents the mean value, bias and mean squared error of the point estimate, respectively. SE represents mean value of the standard error estimate of the slope/slope change; ECP represents the empirical coverage probability of the 95% confidence interval estimate.

| **Table S2**: Simulation results for Scenario 2: synthetic dataset with outliers. Shown are comparisons of point and CI estimation of slope $\xi_{0}$and change of slopes $\xi_{1}, \xi_{2}$ among different statistical methods. | | | | | |
| --- | --- | --- | --- | --- | --- |
| $\xi_{0}=1$ | | | | | |
|  | Est | Bias | MSE | SE | ECP |
| LIMBARE | 0.9955 | -0.0045 | 0.0192 | 0.0326 | 0.994 |
| SMM | -2.6438 | -3.6438 | 10919 | 1.7004 | 0.645 |
| RMESR | 0.8756 | -0.1244 | 4.8986 | 0.0111 | 0.617 |
| CROSS | 0.7891 | -0.2109 | 532.34 | 3.5679 | 0.434 |
| $\xi_{1}=2$ | | | | | |
|  | Est | Bias | MSE | SE | ECP |
| LIMBARE | 2.0049 | 0.0049 | 0.0155 | 0.0348 | 0.998 |
| SMM | 5.4180 | 3.4180 | 10931 | 2.2049 | 0.709 |
| RMESR | 2.1202 | 0.1202 | 4.7873 | 0.0117 | 0.646 |
| CROSS | 0.6163 | -1.3837 | 1628.5 | 1601.4 | 0.857 |
| $\xi_{2}=3$ | | | | | |
|  | Est | Bias | MSE | SE | ECP |
| LIMBARE | 2.9560 | -0.0440 | 0.4241 | 0.1423 | 0.995 |
| SMM | 2.2849 | -0.7151 | 26.872 | 0.8958 | 0.472 |
| RMESR | 2.9496 | -0.0504 | 0.9834 | 0.0150 | 0.578 |
| CROSS | -2.6384 | -5.6384 | 12716 | 1609.7 | 0.788 |

Est, bias and MSE represents the mean value, bias and mean squared error of the point estimate, respectively. SE represents mean value of the standard error estimate of the slope/slope change; ECP represents the empirical coverage probability of the 95% confidence interval estimate.
